# Nocturnal Navigation via a Time-Compensated Lunar Compass in Bull Ants

**DOI:** 10.1101/2025.08.28.672594

**Authors:** Cody A Freas, Ken Cheng

## Abstract

Diurnal navigators are well known to orient via the sun, compensating for its movement over time by linking solar position and speed with terrestrial reference points (Beetz & el Jundi, 2018; Dingle, 2014; Dyer, 1987; Mouritsen & Frost, 2002; Stalleicken et al., 2005; Wehner & Lanfranconi, 1981; Wehner & Müller, 1993). In contrast to the sun, the moon is not only dimmer, but its presence is also highly variable, both temporally and in its illuminance, making it a more complex cue to predict. Yet nocturnal animals employ a celestial compass to orient at night (i.e. the moon and stars; (Dacke et al., 2003, 2013; Dreyer et al., 2025), but navigation via a time-compensated lunar compass has yet to be demonstrated. Nocturnal bull ants (*Myrmecia midas*) navigate home overnight under the dimmest of visual conditions, relying in part on a nocturnal path-integration system informed by lunar cues (Freas et al., 2024). Here, we show the first evidence of a time-compensated lunar compass in a homing animal, with path-integrating ants adjusting path headings by predicting the lunar azimuthal position throughout the night. Ants predict the moon’s future position based upon a generalised internal estimate of the lunar ephemeris. Rather than detailed knowledge of the moon’s nightly schedule, these ants form a lunar path representation via a speed-step function, reflecting the moon’s accelerating then decelerating arc. However, these ants also predictably under- or overestimate the shape of this arc based on individual experience, suggesting a level of temporal uncertainty regarding when the moon rapidly shifts from the eastern to western sky. Additionally, updating this lunar compass relies on the presence of a directionally informative skyline, which allows occasional cross-referencing of lunar movement with external fixed compass cues near the horizon.

## Background

Animal navigators are well known to rely on the sun to maintain correct orientation and efficient homing (Beetz & el Jundi, 2018; Dingle, 2014; Freas & Cheng, 2022; Mouritsen & Frost, 2002; Schmidt-Koenig et al., 1991; Spiecker et al., 2022; Wehner, 2020). The use of a sun compass is especially important for long distance migratory animals which must maintain a specific compass heading over time (Dingle, 2014), but also for the diurnal path integrator to maintain a correct estimate of the nest location (Wehner & Lanfranconi, 1981). As celestial bodies move across the sky, updating their directional position requires compensation of this movement in relation to the goal heading, an ability called a ‘time compensated compass’. The most well studied of these celestial bodies is the movement of the sun, which while its angular speed remains a constant (∼15°/h), its azimuthal position in the sky is more variable, accelerating as it rises, reaching its apex before slowing down as it descends (Wehner & Müller, 1993). Additional complexity arises across days with changes in solar schedule and arc structure, which are dependent upon both the time of year and the navigator’s latitude. This predictable path creates solar ephemeris functions of the sun’s azimuthal direction across each day.

Animal navigators, including insects, are well known to be able to account for the sun’s movement over time, linking temporal periods via an internal clock to the sun’s movement (Jander, 1957; Wehner & Lanfranconi, 1981; Wehner & Müller, 1993). This solar time compensation requires animals to use an external reference point, such as surrounding landmarks, to accurately update (Dyer & Gould, 1981; Towne, 2008). That animal navigators would innately possess detailed representations of these solar ephemeris functions across the year, representative of their specific latitude is unlikely (Wehner, 2020); instead, these navigators employ a variety of simpler, yet highly accurate compensation models to reconstruct a predicted solar ephemeris function based upon experience (Massy & Wotton, 2023). Thus, the general pattern of the sun’s daily course can be approximated for accurate navigation without detailed neural representations of celestial ephemeris functions. For example, honeybees (Dyer & Dickinson, 1994) and desert ants (Wehner & Müller, 1993) possess an innate approximation of the solar ephemeris in the form of a speed-step-function predicting a solar ephemeris in which the sun moves slowly in the morning/evening but rapidly jumps at midday across the south from the eastern to western sky.

We know much about how diurnal animals track the sun, yet many animals successfully orient and navigate at night, when these cues are absent. Some nocturnal animals possess a nighttime sky compass using the position of the moon, the lunar polarisation pattern of the sky or stellar cues to guide their movement (Dacke et al., 2003, 2013; Dreyer et al., 2025; Freas et al., 2024; Klotz & Reid, 1993; Perez et al., 1997; Ugolini et al., 2013; Warrant & Dacke, 2016). Yet, predicting the moon’s position through time is even more complicated than predicting the sun’s, given the moon’s high variability, with the lunar rise schedule, phase, and arc all changing dramatically over the lunar month. This variability makes the possibility of nocturnal animals plotting lunar ephemeris functions difficult to ascertain. To date only sandhoppers have shown evidence of an innate lunar compass, allowing orientation for taxis to or away from the ocean (Ugolini et al., 1999, 2013), yet not navigation.

The large-eyed nocturnal bull ant, *Myrmecia midas*, relies on both solar (during twilight) and lunar cues (overnight) to navigate, as well as learning the terrestrial cues of their established foraging routes (Freas et al., 2018; Freas, Narendra, & Cheng, 2017; Freas, Narendra, Lemesle, et al., 2017; Islam et al., 2020). Outbound navigation occurs during evening twilight, when foragers leave the nest, traveling up to 25m along the forest floor before climbing into the tree canopy to feed (Freas et al., 2018; Freas, Narendra, & Cheng, 2017). Traditionally, these ants have been believed to wait until morning twilight, when light levels begin to rise, before descending their tree and navigating home. Yet, we now know many foragers return to the nest overnight (Freas et al., 2024; Freas, Narendra, & Cheng, 2017). At the extreme, observed during the winter when the night can extend up to 12h, all foragers may return prior to morning twilight (Figure 1A). Such overnight navigation suggests a nocturnal path integrator using celestial compass cues to navigate. These path integrator estimates need to be maintained over many hours (Figure 1A,B) while the ants feed in the canopy, meaning that compensating for lunar movement during these periods would be crucial for keeping an updated sky compass. Recently, it was shown that these nocturnal bull ants can rely on the extremely dim polarisation pattern produced by the moon as part of their path integrator to navigate overnight (Freas et al., 2024; Freas & Cheng, 2025). This pattern acts as a proxy for the moon’s position as it moves through the night, allowing foragers to estimate lunar position even when the moon itself is not visible. Under waning moon conditions, however, when moonrise occurs well after twilight, ants may experience extended cue gaps with no solar or lunar references. Such gaps likely degrade their path integrator, and their reliance on the lunar polarization pattern for homing drops accordingly (Freas et al., 2024). These findings suggest that M. midas may form a short-term, “within-night” time-compensated lunar compass to support overnight navigation. Here, we show that these nocturnal navigators possess an internal time-compensated lunar compass that predicts the moon’s future azimuthal position to guide their navigation, though they show systematic under- and overestimates around the moon’s speed changes at its arc’s apex (Figure 1B). Updating this compass requires reference to a directionally informative skyline, as ants collected before moonrise could not predict later lunar positions without cross-checking against terrestrial cues.

**Figure 1.**
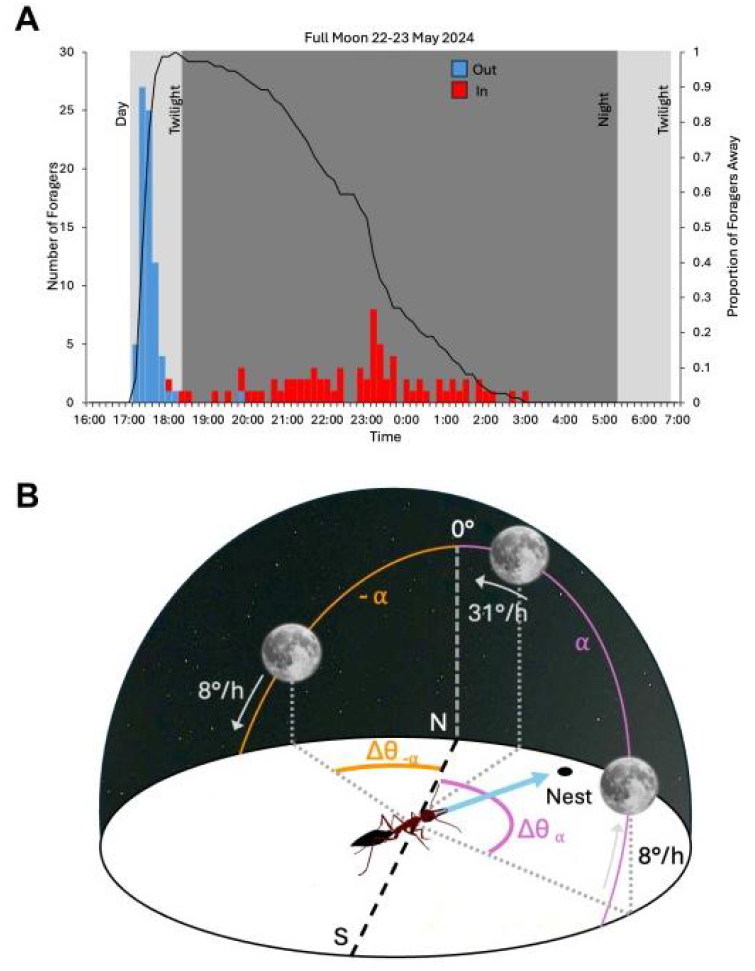
Nightly foraging activity pattern of *M. midas* foragers, showcasing their overnight inbound homing prior to morning twilight. **(A)** Foragers were recorded on the night of May 22-23, 2024, on a full moon night. During this night’s observations, all foragers returned to the nest before the morning twilight, requiring inbound homing when no solar cues were present. This observation is consistent with the foraging ecology of multiple nests, especially in the Australian late fall/winter. Bars, in 10min bins, indicate outgoing foragers leaving the nest area (blue, n=73) and incoming foragers entering the nest area (red, n=73). The black line indicates the overnight foraging activity through time with the proportion of foragers away from the nest throughout the night. **(B)** Diagram of the lunar arc above a navigating *M. midas* using its celestial compass to home to the nest. The moon’s azimuthal speed changes across its arc, beginning slow as it rises in the eastern sky (8°/h). During this rise, the moon’s azimuthal speed accelerates (pink) as it reaches its apex (31°/h) and as it passes 0° in the northern sky. After passing its zenith, the moon begins to slow (orange) again (8°/h) as it sets in the western sky. As *M. midas* foragers begin foraging at twilight and can remain on a single foraging trip for 7h+, maintaining a celestial compass-based path integrator (arrow) should involve compensating for these speed changes.

## Results & Discussion

### Ants predict the lunar arc, but err based on last observed lunar speed

To explore how these ants attend to the moon’s position and account for its movement across the night sky, we tracked the navigational headings of *M. midas* foragers relying solely on their nocturnal path integrator, testing them at a distant site where terrestrial cues were unfamiliar. Foragers from Nest 1 left their nest in the evening twilight on nights while the full moon was visible, accumulating large 14m path-integration-derived vectors. These ants were collected and held either with a clear view of the moon’s movement (*Control*) or were blocked from observing the moon while its horizontal speed either accelerated (*Accelerating Moon*) or decelerated (*Slowing Moon*) across its arc (Figure 1B; Supplemental Figure 1 for Skydomes).

*Control* foragers, with a full view of the night sky over the hold period, were correct in directionally updating their path integrator as the rising full moon accelerated in azimuthal speed moving across the sky. Path headings were correctly oriented in the vector compass direction, being both significantly oriented (Rayleigh Test; Z=7.63; p<0.001) and with the true vector direction (0°) within the 95% Confidence Interval (95% CI) of forager headings (2.8±24.6°; Figure 2A). Under a separate *moonless control*, foragers (n=12) were collected, held and tested identically to *control* foragers, but on a night with the moon well below the horizon (new moon) during the overnight period, providing no visible celestial cues. Here, foragers did not orient at the unfamiliar site, instead exhibiting a uniform distribution of path headings at 2m from release (Rayleigh Test; Z=0.141; p=0.794). The inability to orient to the path integrator overnight when the moon was absent is suggestive that the moon and its associated polarisation pattern are critical parts of the nocturnal celestial compass used by the ants when navigating overnight, at least at this site. Given these nests were located within the greater Sydney area (Macquarie Park, NSW), urban skyglow could make navigation via other celestial cues (i.e., the Milky Way) difficult, whereas in more remote habitats, nocturnal ants may have access to a more robust celestial cue array in the night sky. Thus, the heading changes we observed across conditions can be attributed to foragers orienting via lunar position, since they are unable to orient when the moon is absent.

**Figure 2.**
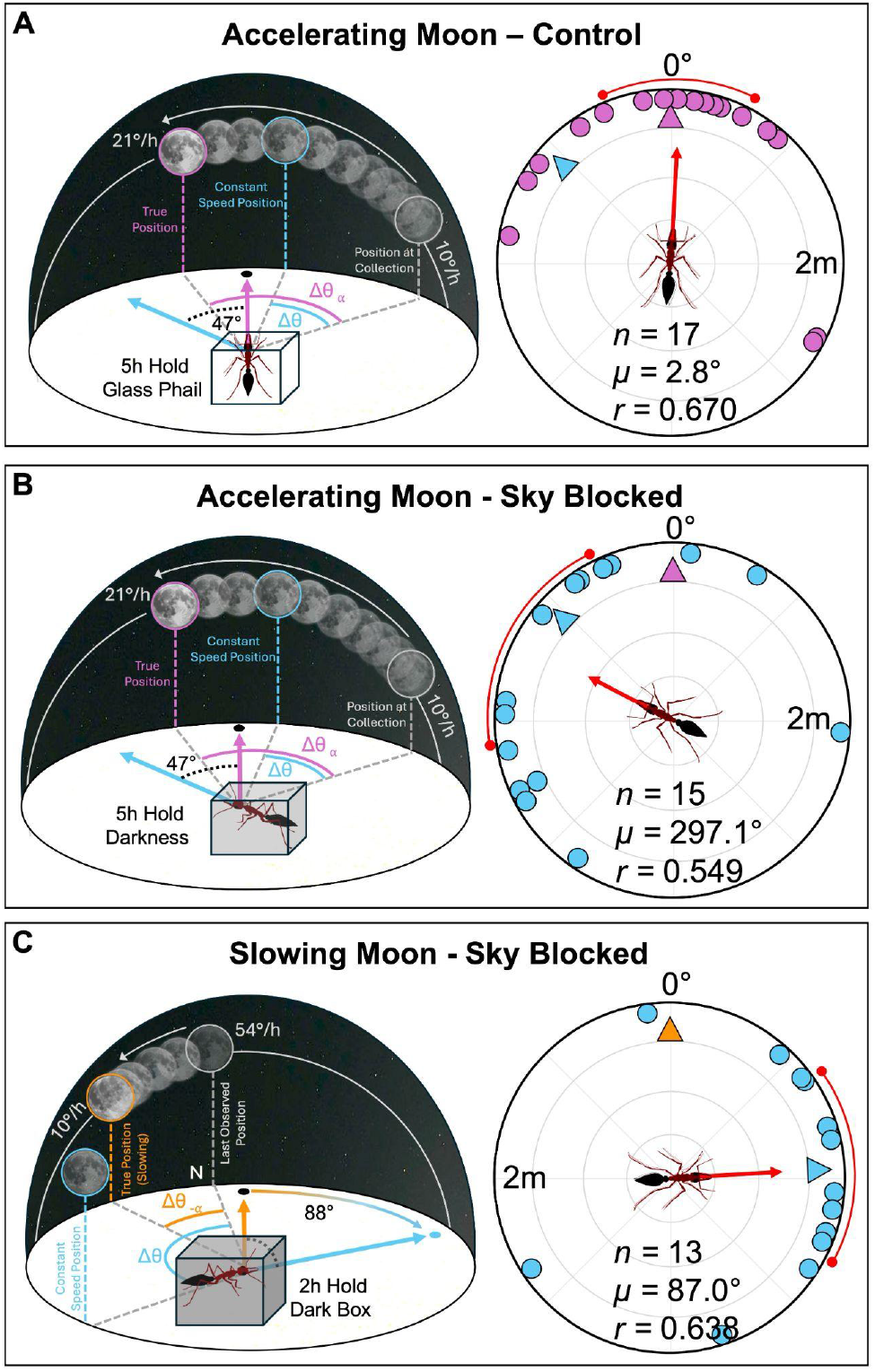
Diagrams and Circular Plots of Forager Headings, at 2m from release, when either allowed or blocked from observing an accelerating or slowing moon. Colours represent different lunar predictions and the resulting predicted vector directions; with blue representing linear extrapolation of lunar azimuthal speed from last observation at collection, while orange and pink represent the true acceleration and deceleration, respectively, of the lunar ephemeris. Nest 1 foragers were allowed to collect a 14m vector to their foraging tree under an accelerating or slowing moon, then either (**A**) held in glass vial and allowed to observe the variable azimuthal speed of the lunar arc or held in darkness while the moon (**B**) accelerated to its apex or (**C**) decelerated during its descent and were then tested at an unfamiliar site. Red arrows within circular plots denote the length (*r*) and direction (µ) of the mean vector, while the red arc represents the 95% CI. n, number of individuals. 0° represents the true vector direction.

In contrast to the controls, *Accelerating Moon* foragers collected under a slow-azimuthal-speed full moon, then held without access to the sky, and released just after the lunar apex (high azimuthal speed) exhibited paths which remained directed (Rayleigh Test; Z=4.53; p=0.009) but with an anticlockwise bias to the left of the true vector direction (0°), with 0° outside the 95% CI of headings (297.1±35.5°; Figure 2B), indicating an underestimation of predicted lunar azimuthal speed over the hold period.

When *Slowing Moon* foragers accumulated their path integrator while the moon (waxing quarter) was moving fast (near lunar apex), held in darkness, and released under a slow-moving moon, foragers exhibited paths showing significant orientation (Rayleigh Test; Z=5.30; p=0.003). However, these headings exhibited a clockwise bias to the right of the true vector direction (0°), with 0° outside the 95% CI of headings (87.0±31.1°; Figure 2C), indicating an overestimation of lunar azimuthal speed. Additionally, both darkness conditions had mean vector directions which were significantly different from *Control* foragers (Control vs. *Accelerating Moon*: Watson-Williams F_(1,30)_=9.18; p=0.005; *α*=0.05; Control vs. *Slowing Moon*: Watson-Williams F_(1,28)_=16.24; p<0.001; *α*=0.025). Heading variance did not significantly differ between conditions (Var Tests; p > 0.05).

Results indicate predictable under/over estimations of future lunar azimuthal positions based upon the moon’s movement when it was last observed during vector accumulation. These heading biases are generally in line with work on the solar ephemeris function in desert ants, with similar under/overestimated predictions of solar positioning (Wehner, 2020; Wehner & Lanfranconi, 1981). In the current study, however, prediction errors of lunar position were much more pronounced. *M. midas* headings aligned well with the hypothesis that nocturnal ants predict lunar movement by extrapolating a constant lunar speed based on the last observed speed at collection. Vector orientation using this predicted lunar position (constant lunar speed from last observation) was within the 95% CI of headings in both conditions (Figure 2B,C, blue arrows). This contrasts with diurnal ants, which while exhibiting prediction errors when held for short periods (1h), also show foragers correctly estimating the solar speed-step during midday (Wehner & Lanfranconi, 1981; Wehner & Müller, 1993). The solar midday and its corresponding speed-step across days is quite stable compared to the moon, for which speed steps (if measured by lunar apex) are ∼40–60min later each subsequent night. Given this high lunar variability over even a handful of days, and that the current tests have collection or tests periods which occurred during to the lunar speed-step (See Figure 3; Supplemental Figure 2), these tests cannot, in isolation, differentiate between this linear extrapolation model and a more complex lunar ephemeris-function prediction system within *M. midas’* lunar compass. In particular, foragers may struggle to predict the exact timing of the lunar speed-step function.

**Figure 3.**
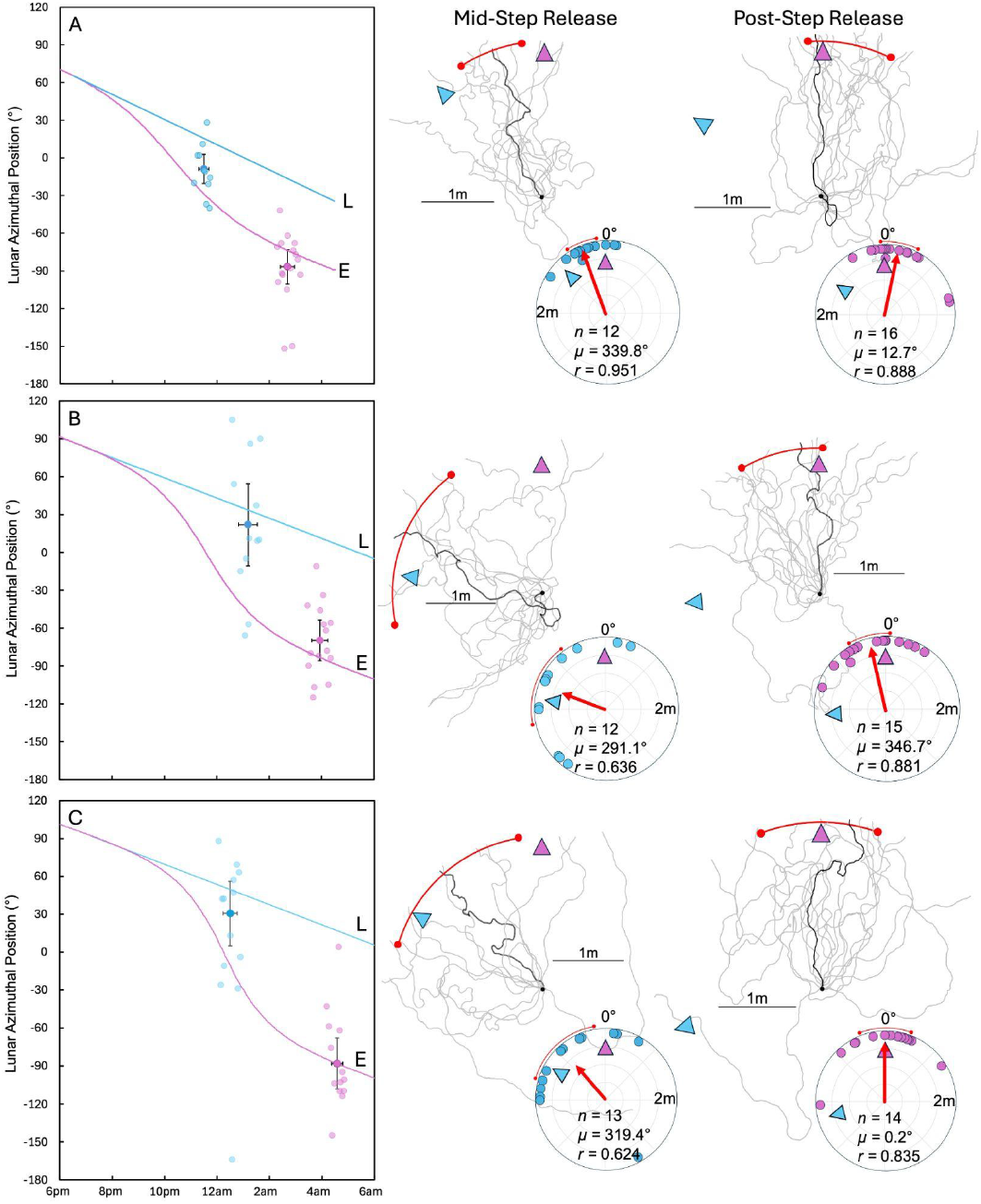
Lunar Ephemeris Functions, Forager Paths and Circular Plots of Conditions and Forager Headings, at 2m from release. Foragers were blocked from observing an accelerating moon after collection (black circle) and released either during the lunar 0° step (Mid-Step) or after the lunar 0° step (Post-Step). (**A**) Nest 1, April 10, 2025; (**B**) Nest 2, April 12, 2025; (**C**) Nest 3, April 13, 2025. The blue line represents the lunar azimuthal position based upon linear extrapolation from the last observed speed at collection (L) while the pink line represents the true lunar ephemeris function (E). Individual forager headings are plotted along the true lunar position at test time (line E), with alignment to this line indicating orientation to the true path integrator direction. Error bars represent the 95% CI of headings (x axis) and the standard deviation of mean test time. Red arrows within circular plots denote the length (*r*) and direction (µ) of the mean vector, while the red arc represents the 95% CI. n, number of individuals. 0° represents the true vector direction.

**Figure 4.**
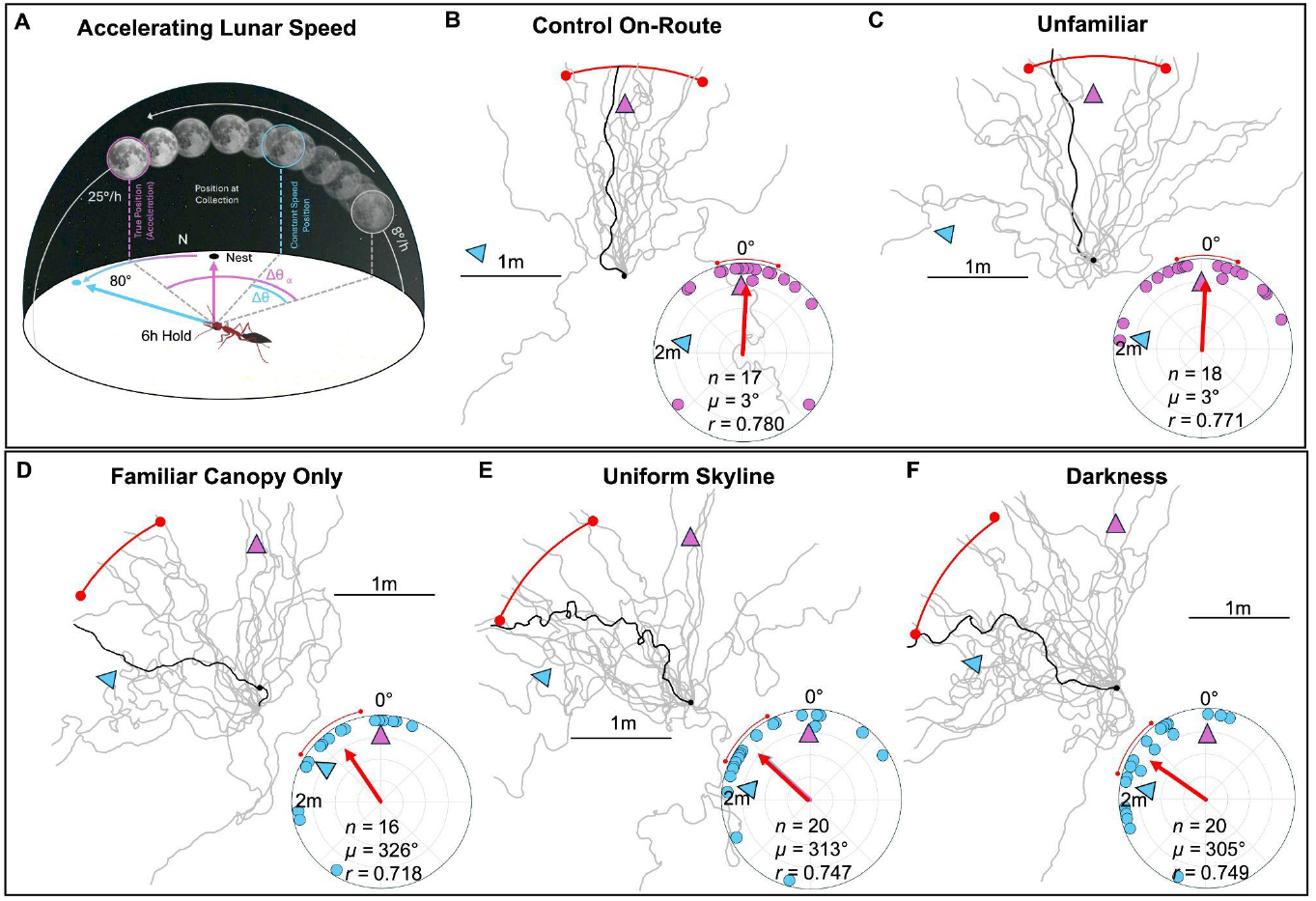
Experimental Diagram, Forager Paths and Circular Plots of Conditions and Forager Headings, at 2m from release. (**A**) Diagram of the lunar arc’s acceleration from collection and its effect on the ant’s predicted vector direction. (**B**) Control foragers were held in a glass vial while along the known route. (**C**) Unfamiliar foragers were held at a distant unfamiliar site, distinct from the testing site. (**D**) Canopy foragers were allowed to see the sky and overhead canopy but were blocked from observing the skyline below 45°. (**E**) *Uniform Skyline* foragers were allowed to see only the overhead sky while blocked from observing the skyline below 45°. (**F**) *Darkness* foragers were held in darkness during the hold period. Within the condition diagrams, colours represent different lunar predictions and the resulting predicted vector directions; with blue representing the vector direction based upon linear extrapolation prediction of lunar azimuthal speed from last observation at collection under a rising moon, while pink represents the true acceleration of the lunar ephemeris. Red arrows within circular plots denote the length (*r*) and direction (µ) of the mean vector, while the red arc represents the 95% CI. n, number of individuals. 0° represents the true vector direction.

### Linear extrapolation or lunar ephemeris function?

We next investigated whether nocturnal ants predict the moon’s movement using simple linear extrapolation or a more complex lunar ephemeris function. At three nests, outbound foragers were collected at twilight, when the moon was rising in the east and moving at low azimuthal speed. Foragers were held in darkness and then tested during one of two distinct lunar phases: (1) the ‘mid-step’, when azimuthal speed peaked near the lunar apex, or (2)’post-step’, when the moon’s speed had slowed and its position shifted from the eastern to the western sky. Importantly, predictions based on linear extrapolation and a lunar ephemeris function directionally diverge during the lunar speed-step, with this divergence being maintained as the moon slows during its descent (Figure 3, blue vs. pink lines).

At all three nests, foragers tested mid-step, near the lunar apex, underestimated the amount of lunar azimuthal change, resulting in path headings shifted in the anticlockwise direction away from the true vector (Figure 3). At all three nests, foragers were well oriented (Rayleigh Test; p<0.001), but in each the true vector direction (0°) was to the right of the 95% CI of headings (Nest 1: 339.8±11.8°, Figure 3A; Nest 2: 291.1±32.5°, Figure 3B; Nest 3: 319.42±32.0°, Figure 3C). At Nest 1, the mean vector direction of headings exhibited a midpoint between the true lunar ephemeris and an extrapolated constant lunar speed from collection (Both 316° and 0° were outside 95% CI; Figure 3A). At Nests 2 and 3, in contrast, heading directions were in line with foragers extrapolating lunar speed linearly, with this direction (Nest 2: 279°; Nest 3: 300°) within the 95% CI of headings.

This underestimation of lunar speed did not persist post the speed-step, as foragers tested later in the same night showed a step-down shift in headings aligning with the true lunar ephemeris, showcased through orientation in the vector compass direction (Figure 3A-C). Foragers held in darkness post the lunar apex when the descending moon’s azimuthal speed had returned to near that at ascending collection (with the moon now in the western sky) deviated significantly from the linear extrapolation prediction, ‘L’ (Figure 3, Blue line) and were well aligned with an accurate prediction of the lunar position via an ephemeris function over the delay (Figure 3, ‘E’, Pink line). Headings were significantly oriented at all nests (Rayleigh Test; p < 0.001) and headings were well oriented to their inbound vector direction (0°), being within the 95% CI of headings at all three nests (Nest 1: 12.7±13.7°; Nest 2: 346.7±16.2°; Nest 3: 0.2±20.1°). At all nests, mean heading directions significantly differed between Mid-Step and Post-Step release foragers (Watson-Willaims test; Nest 1: F_(1,26)_=10.371; p=0.002; Nest 2: F_(1,25)_=10.371; p=0.004; Nest 3: F_(1,25)_=4.856; p=0.037). While heading variance was higher during mid-step testing at Nest 2 and Nest 3 (Figure 3B,C), Var Tests indicated that these increases were not significant (p > 0.05).

Foragers’ accurate headings post-step were indicative that these foragers have incorporated at least the general shape of the lunar ephemeris function into their neural compass system: the full moon takes position in the east during the first part of the night before rapidly shifting to the western sky later in the night. This pattern is strikingly akin to the solar ephemeris function observed in both desert ants and honeybees tested in the morning and afternoon (Dyer & Dickinson, 1994; Wehner, 2020; Wehner & Lanfranconi, 1981), with both ants and bees predicting a speed-step-function near mid-day. Here, we find a pattern consistent with the ants predicting a speed-step function across the northern sky as the moon rapidly moves near its apex from the east to west. Thus, these ants are accurate at predicting the moon’s position after the step, but predicting lunar positioning during the step appears more difficult, with most of our evidence suggesting a majority of foragers are aligning with linear extrapolation (except mid-step foragers at Nest 1, Figure 3A). This may be indicative of how lunar movement predictions are tied to experience outside of the step. Given the lunar arc’s complexity exceeds the sun’s, it is unlikely that ants possess a highly detailed and accurate lunar ephemeris function across nights. They are more likely to use a simpler solution based upon the general characteristics of the lunar arc (Massy & Wotton, 2023). This solution would predict that the moon would rise in the east at a relatively constant slow azimuthal speed, possibly relying on observed speeds to predict pre-step lunar movement, before quickly shifting to the western sky where its speed would again slow. Thus, foragers would perform quite well at orienting both during lunar rise (linear extrapolation) and once the step from east to west has been predicted (linear extrapolation + speed-step). During the step period, however, predictions appear to become less accurate and are influenced by the last observed speed. Mid-step tested foragers show evidence of uncertainty over whether the lunar speed-step has occurred, with some foragers orienting correctly while the majority clearly have not, remaining oriented in accordance with the pre-step lunar azimuthal speed based upon linear extrapolation of their last lunar observation. Testing for bimodality in the data is difficult with this number of foragers; however, a similar pattern of uncertainty near the solar apex is present in honeybees (Dyer & Dickinson, 1994), see their Figure 1), suggesting that the internal timing of the step function in both the sun and moon may be imprecise. Both patterns of data suggest that similar rule-of-thumb prediction processes may underlie tracking both solar and lunar ephemeris in Hymenoptera.

Findings suggest the internal representation of the *M. midas* time-compensated lunar compass incorporates a linear extrapolation of lunar speed during easterly ascent and westerly descent with a speed-step acceleration then deceleration during lunar apex in the northern sky (Supplemental Figure 2). That foragers are well oriented under a descending moon suggests the speed-step has been predicted, while the consistent anticlockwise bias during the speed-step suggests foragers predict a linear rate of movement while the moon is ascending. That the lunar acceleration period near its apex is less predictable may result from the variability of this temporal period across nights. For example, across the three nights of April testing (night of 10^th^ to 13^th^), lunar apex shifted from 22:16 on to 00:15 (just after midnight on the 14th), a 119min difference over only a few days. Contrast this with the relative stability of solar noon, which only changes by 1min between these dates. Given this high variability, the lunar arc may be hard to predict, with foragers delaying when they expect the speed-step to occur, resulting in the observed bias in headings. Importantly, these foragers are likely highly experienced with the foraging route on previous nights, with individuals being long-lived (1 year) and foraging most nights, suggesting there may be some lunar-apex delay prediction built into this internal representation from one night to the next.

### A within-night prediction mechanism

Given the moon’s variability across the lunar month, a detailed ephemeris across nights appears unlikely, especially given foragers’ temporal uncertainty regarding the speed-step. However, since *Myrmecia* foragers are long-lived and forage almost nightly, they might be predicted to retain memories from previous nights, maintaining longer-term lunar predictions. To test whether foragers can orient using such long-term predictions when first encountering the moon on a given night, we collected outbound foragers at twilight, when no lunar cues were visible (waning quarter moon, ∼50% illuminance). We held them in darkness until after moonrise and then released them at the distant, unfamiliar site once the quarter moon was visible. These foragers did not orient significantly, instead showing random headings at 2m from release (Rayleigh test: z = 0.30, p = 0.748).

These random orientations are consistent with lunar prediction being a within-night process. Foragers appear to begin a predictive azimuthal speed calculation only once they first observe the moon, though it is unknown if this is possible when the moon rises later in the night, well after twilight. This interpretation is consistent with previous work showing that if the moon is absent for part of the night, ants rely less on its polarization pattern (lunar position proxy) than if it is continuously visible (Freas et al., 2024). Thus, a potential cue gap in the sky (with the moon absent at the end of solar twilight) may degrade the overnight path integrator. Although we cannot systematically reconstruct each forager’s experience on previous nights, they likely foraged on one or more of these nights as well. Nevertheless, any memory of the lunar arc from prior nights was insufficient for orientation on the test night. Whether lunar predictions are strictly within-night, as these results suggest, or whether some longer-term predictions are possible, remains an open question.

The lack of orientation in these headings also suggests that this species neither exhibits phototaxis towards the moon’s position nor orient to the lunar position as if it were the sun, as is the case with diurnal ants attempting to orient at night (Lanfranconi, 1982). Whether these nocturnal ants possess a solar ephemeris despite only foraging after sunset is unknown.

### Terrestrial Cues near the skyline critical for lunar compass compensation overnight

As foragers with a view of the moon overnight accurately counteract the observed prediction errors under variable lunar speeds, updating the lunar ephemeris, especially around the step, likely requires some form of earthbound reference to occasionally update lunar changes. The skyline panorama is a likely candidate, as it is both known to be used for solar azimuthal tracking in bees (Dyer & Gould, 1981; Towne, 2008; Towne & Moscrip, 2008) and is the most critical terrestrial cue that these nocturnal ants attend to (Islam et al., 2022).

To assess what external cues are used to track lunar movement, foragers which had accumulated a large 23m path integrator (Nest 3) under a slow rising moon were held and then tested near the lunar apex (∼45° elevation), with individuals assigned to one of five conditions during captivity with different levels of visual information present: 1.) *Control* foragers were allowed a full view of the skyline and overhead sky, held within a glass vial, at a known mid-point along their route; 2.) *Unfamiliar* foragers were allowed a full view of the skyline and overhead sky at a distant unfamiliar location; 3.) *Uniform Skyline* foragers were allowed a full view of the sky above 45° elevation, with the view below 45° blocked with a uniform-skyline arena; 4.) *Canopy* foragers were held in identical uniform-skyline arenas but on-route beneath a familiar overhead canopy with access to the sky; 5.) *Darkness* foragers were held in darkness with all visual cues blocked.

Under an accelerating moon, *Control* foragers, exhibited paths which were significantly oriented (Rayleigh Test; Z = 10.350; p < 0.001) and in the vector direction, with the true vector compass direction (0°) well within the 95% CI of headings (3.4±19.0°). Paths of *unfamiliar* foragers were also significantly oriented in the true vector direction (Rayleigh Test; Z = 10.690; p < 0.001), with the true vector compass direction (0°) within the 95% CI of headings (2.5±18.9°). The mean vector direction of *unfamiliar* and *Control* foragers (2.5° vs. 3.4°) did not significantly differ (Watson-Williams; F_(1,33)_ = 0.004; p = 0.948), suggesting skyline familiarity did not influence lunar compass updating. *Canopy* foragers were significantly oriented (Rayleigh Test; Z = 8.244; p < 0.001), but these paths were biased anticlockwise of the true vector direction (0°), with 0° outside of the 95% CI of headings (324.8±22.8°). The mean vector direction of *Canopy* forager paths (324.8°) was significantly to the left of *Control* foragers (F_(1,31)_ = 6.029; p = 0.02; *α* = 0.025). Foragers held in a *Uniform* skyline with no canopy were significantly oriented (Rayleigh Test; Z = 11.167; p < 0.001), but again paths exhibited an anticlockwise bias to the left of the true vector direction (0°), with 0° outside the 95% CI of headings (313.4±19.0°) and with the mean vector direction significantly to the left of *Control* foragers (F_(1,35)_ = 12.084; p = 0.001; *α* = 0.017). Foragers held in *Darkness* were significantly oriented (Rayleigh Test; Z = 11.223; p < 0.001), and again forager paths exhibited an anticlockwise bias to the left of the true vector direction (0°), with 0° outside the 95% CI of headings (305.2±19.2°), and the mean vector direction of this condition was again significantly to the left of *Control* foragers (F_(1,35)_ = 16.389; p < 0.001; *α* = 0.013). In all three conditions where an anticlockwise bias was observed, ant headings also did not align with pure linear extrapolation of lunar speed, with both the true vector direction (variable lunar speed, Figure 3C-E, Pink) and the constant-lunar-speed extrapolation (Figure 3C-E, Blue) outside the 95% CI of headings.

Two of these conditions (*Control* and *Uniform Skyline*) were repeated under a slowing moon, with ants collected near the lunar apex and tested under a setting moon (Supplemental Figure 3). Under a slowing moon, *Control* foragers with a full view of the night sky during holding exhibited paths which were significantly oriented (Rayleigh Test; Z = 7.756; p < 0.001) and the true vector compass direction (0°) was well within the 95% CI of headings (348.1±27.4°). In contrast, foragers held in a *Uniform* skyline with a clear view of the sky (Supplemental Figure 3), were significantly oriented (Rayleigh Test; Z = 7.522; p < 0.001), but paths exhibited a clockwise bias to the right of the true vector direction (0°), with 0° outside the 95% CI of headings (58.3±27.5°), with the mean vector direction significantly to the right of *Control* foragers (F_(1,25)_ = 11.506; p = 0.002). Here, heading directions (95% CI) were in line with linear extrapolation of lunar speed from collection (Supplemental Figure 3).

The observed bias in headings suggests that continual updating of the lunar compass through the night requires not only a view of the night sky, but also occasional cross-referencing with the surrounding terrestrial cues, either familiar or initially unfamiliar, near the skyline. Testing occurred on nights in which the lunar arc travelled above 45°, suggesting that even direct observation of the moon under a directionally uninformative skyline is insufficient to update the lunar compass correctly. When the skyline is unavailable, foragers appear to fall back on their predictions of the general lunar ephemeris, under/overestimating lunar azimuthal movement based upon their last usable observation, before collection. Again, we see the same uncertainty of the lunar path around the lunar apex, suggesting foragers struggle to time the lunar step from the eastern to the western sky. Interestingly, the initial familiarity of this skyline was not critical, as foragers which observed the moon’s movement at an initially unfamiliar site were still able to accurately track the lunar arc and oriented correctly when homing.

## Conclusions

In a first for navigating animals, we show that *M. midas* ants predict aspects of the lunar arc overnight as part of their nocturnal path integrator via a time-compensated lunar compass. The nature of this prediction strategy is consistent with a simplified lunar ephemeris function: slow azimuthal changes at rise and set predicted by linear extrapolation, and a rapid speed-step adjustment near the apex. Given the moon’s high between night temporal variability, ants struggle with predicting the speed-step. Our findings further indicate that lunar prediction in *M. midas* is a within-night process, with foragers initiating lunar position estimates only after first observing the moon each night, rather than relying on memory from prior nights. Finally, we show that updating the lunar compass over extended periods may require occasional cross-referencing with the terrestrial skyline. In the absence of this cue, foragers appear to rely more heavily on predicting the lunar arc. These findings suggest that lunar movement prediction strategies in M. midas share broad similarities with how diurnal Hymenoptera predict the sun’s arc, despite the moon’s greater variability, highlighting the flexibility of the insect path integrator across both day and night.

## Materials and Methods

### Study Site and Species

Experiments were conducted from May 2024 through April 2025 on three *Myrmecia midas* nests on the Macquarie University Wallumattagal campus in Sydney, Australia (33°46 11S, 151°06 40E). Nests were chosen where *M. midas* foragers travelled >10m to a nearby foraging tree (Supplemental Figure 1) to ensure individuals orient to their path integrator upon release (Freas, Narendra, & Cheng, 2017; Freas, Narendra, Lemesle, et al., 2017). Nests typically occur within *Eucalyptus* tree stands, at the base of a tree (Deeti et al., 2024). *M. midas* forages nocturnally, with foraging onset occurring ∼20–40min after sunset when foragers leave the nest to travel to and up one of several surrounding trees to feed overnight (Freas et al., 2018). During this twilight period, solar cues are still present and these ants actively rely on these cues for vector accumulation (Freas, Narendra, & Cheng, 2017; Freas, Narendra, Lemesle, et al., 2017). In contrast, inbound navigation is highly temporally variable, with foragers descending the tree and navigating to the nest either overnight or into morning twilight, meaning many foragers navigate home at night, in the absence of solar cues (Freas et al., 2024; Freas, Narendra, & Cheng, 2017). All collections across tests occurred at the base of individuals’ foraging trees while the moon was clearly visible along the foraging route. As foragers leave the nest individually over a ∼40min period, foragers in each condition were held in 7cm diameter circular glass vials along their foraging route until the start of the holding period. Each ant was then randomly assigned to a condition (i.e. placed within a darkened box) with hold conditions beginning at the same time. This allowed all foragers across conditions to observe the moon’s movement until the moment they were assigned a condition. All individuals were provided a small amount of honey during holding to promote inbound homing during testing. All testing occurred at an unfamiliar site (>200m from all nests), under clear, cloudless nights where the moon was clearly visible to foragers both during collection and testing.

### Lunar Arc and Phase

For all testing conditions involving accelerating lunar speed, evening twilight collection under slow azimuthal speed requires rising moons near full phase (≥90% illuminated), since only then does the lunar cycle place the moon just above the eastern horizon at twilight. Conversely, twilight collection during slowing moons requires the moon to be near its apex, when azimuthal speed is maximized. This alignment occurs around the waxing quarter (∼50% illuminated), when the lunar schedule places it at its apex during evening twilight. For each testing night, the moon’s arc across the horizon was obtained from the Astronomical Almanac (http://asa.usno.navy.mil).

### Accelerating/Slowing Lunar Speed Experiment

Foragers from Nest 1 were placed in two conditions, a *Control* (n=17) and a *Sky Blocked* condition (n=15), both under an accelerating moon (Figure 1A,B; 98% Illumination). Outbound individuals were allowed to accumulate a path integrator (14m) under a slowly moving (10°/h azimuthal) rising moon (20°–30° elevation) and collected near the base of their foraging tree. *Control* foragers were collected and held in glass phials placed along their foraging route while *Sky Blocked* foragers were held within a darkened box at the same location. After the 5h hold period, the moon had risen to near its apex, accelerating to 22°/h (40°–48° elevation), creating a 47° discrepancy between the vector direction under the true variable lunar speed and a linear-extrapolation prediction process. Foragers were transferred to the unfamiliar site and were individually released and allowed to choose a travel heading at the unfamiliar site. At 2m from release, their heading direction was recorded, and each forager was marked as tested and returned to its foraging tree.

On a separate night, foragers (n=13) were collected and held in a darkened box after accumulating their path integrator under a fast-moving moon (*Sky Blocked Slowing Moon*), near its apex (54°/h azimuthal; 74° elevation; 62% Illumination). Hold times for this condition were 2h, over which the moon moved from its apex to the western sky, decelerating to 10°/h (40°–48° elevation). Foragers were then transferred to the unfamiliar site and tested identically to the two previous conditions.

In the separate group of foragers at Nest 1 (n=12) were collected and tested as a *Moonless Control*. These foragers were collected and tested at the unfamiliar site identically to *Control* foragers but on a cloudless, ‘new moon’ night when the moon had a near 0% illuminance, reflecting little to no sunlight. New moons correspond with the moon being above the horizon during the day so for our testing, the moon was also well below (>-20°) the horizon during both path-integrator accumulation (evening twilight) and overnight testing. Headings were collected at 2m at the unfamiliar site.

### Lunar Step Function Experiment

On separate nights, outbound foragers from three nests were allowed to accumulate their path integrator to their foraging tree (Nest 1: 14m; Nest 2: 12m; Nest 3: 23m; Figure 2) under a slowly moving rising moon (Azimuthal speed, Elevation, illumination; Nest 1: 10°/h, 25°, 95%; Nest 2: 8°/h; 23°; 99%; Nest 3: 9°/h, 24°, 100%). At each nest foragers were held within a darkened box, with one condition tested at the unfamiliar site when the moon was near its apex, during the high azimuthal speeds of the Step Function (Nest 1: 23°/h; Nest 2: 29°/h; Nest 3: 40°/h) and another condition tested at the same site, well after the step, as the moon slowed and was setting in the western sky (Nest 1: 11°/h; Nest 2: 10°/h; Nest 3: 10°/h). Upon release, each forager was tracked, and their paths were recorded to 2m with pen and paper and their heading directions at 2m were collected. Tested foragers were marked and returned to their foraging tree.

### Moonless Vector Accumulation

To assess if lunar time compensation in these ants represents a long-term representation of the lunar arc across nights or if predictions occur on shorter-term ‘within-night’ predictions, we collected foragers under a waning quarter moon (∼50% illuminance), corresponding with the moon being absent during vector accumulation (evening twilight), with moonrise occurring in the middle of the night. Here, Nest 1 foragers (n=17) were collected as they reached their foraging tree and placed into a darkened box, blocking them from observing celestial cues until testing. Foragers were released at the unfamiliar site at 1am when the moon was rising in the east and their headings at 2m were recorded.

### Cross Referencing Visual Terrestrial Cues Experiment

At Nest 3, outbound foragers were allowed to accumulate their path integrator to a foraging tree 23m from the nest (under a slow moving moonrise ; 8°/h; 20°–28° elevation; +95% illumination) where they were collected and designated to one of five hold conditions; *Control* foragers (n=17) were held in glass vials along their foraging route with a clear view of the night sky (Figure 3A). *Unfamiliar* foragers (n=18) were held in glass vials at a directionally distinct but unfamiliar site (visually independent of the unfamiliar test site), with a clear view of the sky (Figure 3B; Supplemental Figure 1). *Uniform* Skyline foragers (n=20) were held with a view of the sky above 45° but with the terrestrial scene rendered directionally uninformative, blocked with a uniform arena placed around each vial (Figure 3C). *Canopy* foragers (n=16), also had their skyline rendered uninformative via an uniform arena but were held on-route beneath their familiar foraging tree canopy with access to the sky (Figure 3D). Finally, *Darkness* Foragers (n=20) were held in darkness with all visual cues blocked, identically to previous sky blocked conditions (Figure 3E). Foragers in all conditions were tested during the lunar step near its apex, as the moon had accelerated to 25°/h, a period at which foragers under-estimate lunar azimuthal movement. Upon release, each forager was tracked, and their paths were recorded to 2m with pen and paper and their heading directions at 2m were collected. Tested foragers were marked and returned to their foraging tree.

As a final control test under a slowing moon for the uniform skyline condition, we collected and tested a separate group of individuals under *Contro*l (n=12) and *Uniform Skyline* (n=14) conditions, but instead of foragers accumulating their path integrator under a slow moon, in this group foragers did so under a fast moon near its apex (26°/h; 43° elevation; 80% illumination). Foragers were held until the moon slowed (9°/h; 23° elevation) and tested identically to previous conditions at the unfamiliar test site.

### Statistical Analysis

Data were analysed with circular statistics (Batschelet, 1981). To determine if forager headings shared a common direction, we used Rayleigh tests for circular data (Fisher, 1993). If forager headings were directed, we further analysed whether the mean heading direction was in the vector direction based upon separate lunar movement prediction models (linear extrapolation vs. true lunar ephemeris) using the 95% confidence interval (CI). Between-condition comparisons of 2m headings were conducted using Watson-Williams F-tests with Holm-Bonferroni corrections to the critical p value for multiple comparisons. Watson–Williams tests are sensitive to differences in circular variance; thus when differences between conditions were significant, we further analysed these conditions with Var tests to assess if these differences were due to variance (Wystrach et al., 2014). Var tests consist of calculating the absolute angular error from the mean vector for all the headings in each condition and this error magnitude was compared between conditions using Mann-Whitney U tests.

## Acknowledgements

This work was conducted upon the grounds of Macquarie University whom we thank for access to the nests. We thank Rüdiger Wehner and Jochen Zeil for their comments on a previous version of this manuscript.

## Contributions

C.A.F. - Conceptualization, Data curation, Formal analysis, Funding acquisition, Validation, Investigation, Visualization, Methodology, Writing.

K.C - Funding acquisition, Writing - Revisions.

## Supplemental Figures

**Supplemental Figure 1.**
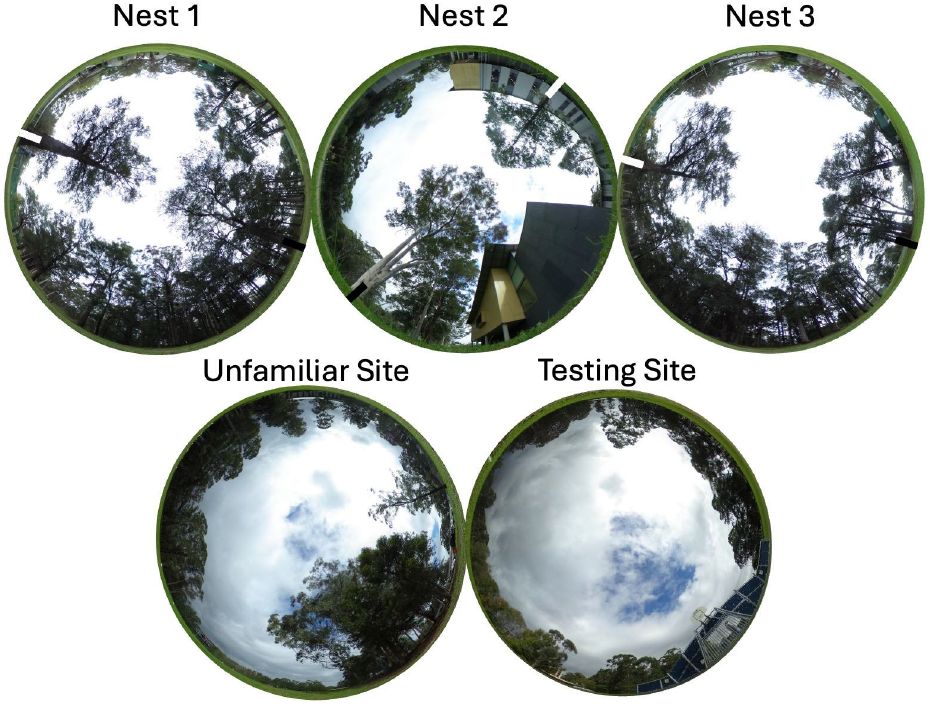
Images of the landmarks, sky and canopy cover at nests and testing sites. Photos of nests were taken at the on-route midpoint between the foraging and nest trees. Black bars represent each nest tree while white bars represent the foraging tree. The unfamiliar site panorama/sky was used for the U*nfamiliar* condition, where foragers were held with a clear view of the moon but at a previously unfamiliar site. The Test site image was where all foragers were tested.

**Supplemental Figure 2.**
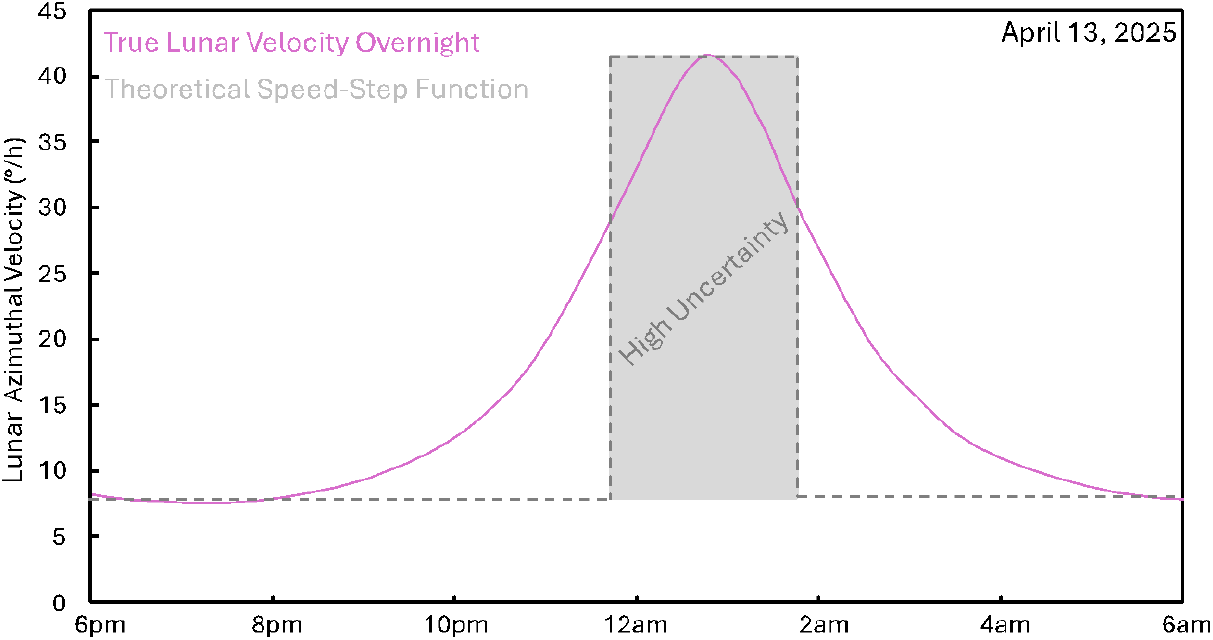
Lunar Speed-Step-Function prediction for the lunar time compensated compass. The true lunar speed spike on the 13^th^ of April 2025 (pink) overlayed with the proposed lunar speed-step-function in the nocturnal path integrator (grey). The grey area represents an area of temporal uncertainty within the system predicting this speed-step, which could explain which foragers underestimate lunar speed during this period, but remain well oriented to the setting moon, post speed-step.

**Supplemental Figure 3.**
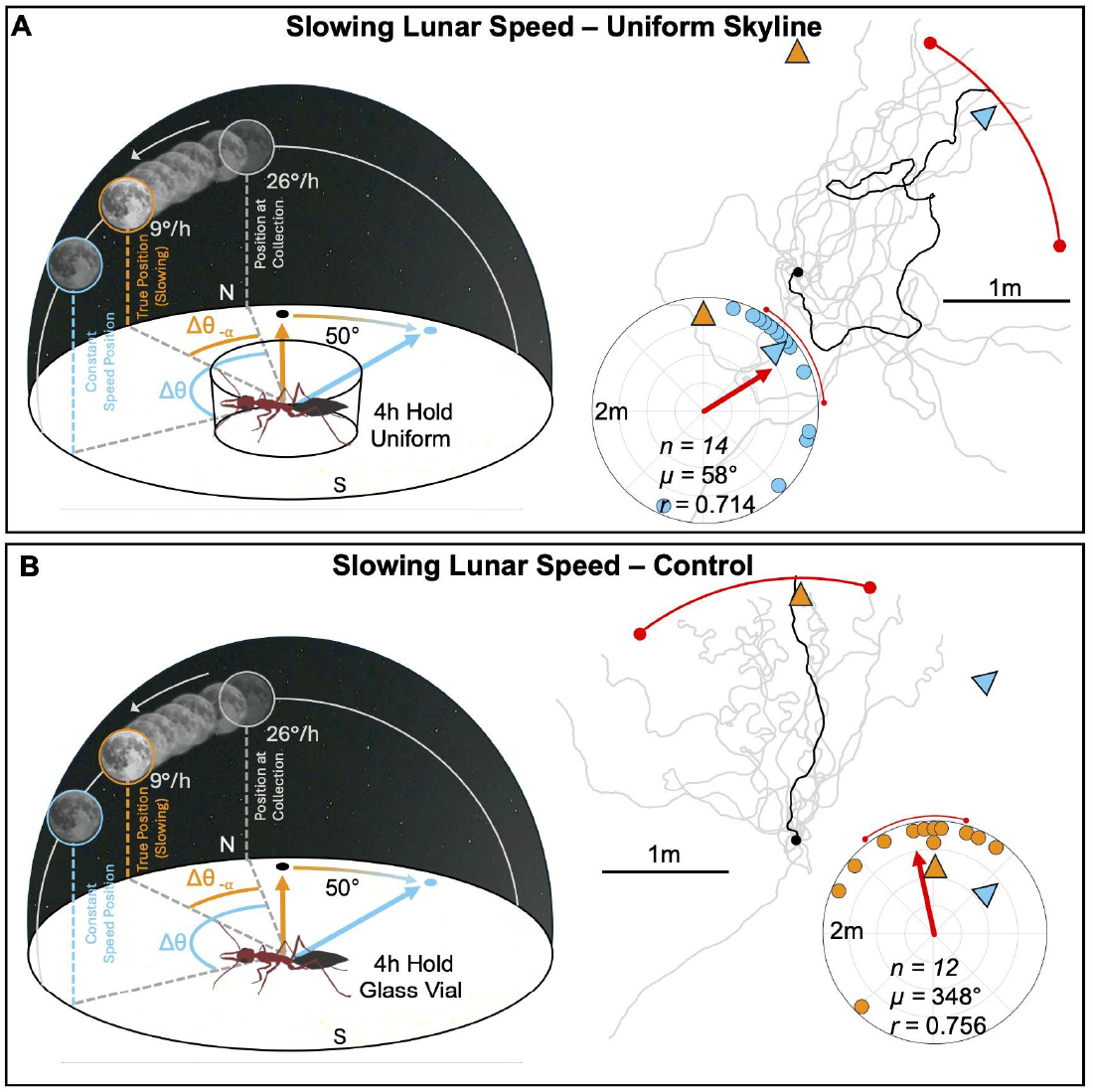
Experimental Diagrams, Forager Paths and Circular Plots of Conditions and Forager Headings, at 2m from release, when allowed to observe a slowing moon with or without access to a directionally informative terrestrial skyline. Within the condition diagrams, colours represent different lunar predictions and the resulting predicted vector directions (triangles); with blue representing linear extrapolation of lunar azimuthal speed from last observation at collection, while orange represents the true deceleration of the lunar ephemeris. Foragers were allowed to collect a path integrator vector to their foraging tree under a fast moving moon near its apex, then either (**A**) held with a uniform skyline or (**B**) held in a glass vial while the moon decelerated during its descent and were then tested at a distant unfamiliar site. Red arrows within circular plots denote the length (*r*) and direction (µ) of the mean vector, while the red arc represents the 95% CI. n, number of individuals. 0° represents the true vector direction.

## References

Batschelet, E. (with Internet Archive). (1981). Circular statistics in biology. London ; New York: Academic Press. http://archive.org/details/circularstatisti0000bats

Beetz, M. J., & el Jundi, B. (2018). Insect Orientation: Stay on Course with the Sun. Current Biology, 28(17), R933–R936. 10.1016/j.cub.2018.07.032

Dacke, M., Baird, E., Byrne, M., Scholtz, C. H., & Warrant, E. J. (2013). Dung Beetles Use the Milky Way for Orientation. Current Biology, 23(4), 298–300. 10.1016/j.cub.2012.12.034

Dacke, M., Nilsson, D.-E., Scholtz, C. H., Byrne, M., & Warrant, E. J. (2003). Insect orientation to polarized moonlight. Nature, 424(6944), 33–33. 10.1038/424033a

Deeti, S., Tjung, I., Freas, C., Murray, T., & Cheng, K. (2024). Heavy rainfall induced colony fission and nest relocation in nocturnal bull ants (Myrmecia midas). Biologia, 79(5), 1439–1450. 10.1007/s11756-024-01634-4

Dingle, H. (2014). Migration: The Biology of Life on the Move. Oxford University Press. 10.1093/acprof:oso/9780199640386.001.0001

Dreyer, D., Adden, A., Chen, H., Frost, B., Mouritsen, H., Xu, J., Green, K., Whitehouse, M., Chahl, J., Wallace, J., Hu, G., Foster, J., Heinze, S., & Warrant, E. (2025). Bogong moths use a stellar compass for long-distance navigation at night. Nature, 643(8073), 994–1000. 10.1038/s41586-025-09135-3

Dyer, F. C. (1987). Memory and sun compensation by honey bees. Journal of Comparative Physiology A, 160(5), 621–633. 10.1007/BF00611935

Dyer, F. C., & Dickinson, J. A. (1994). Development of sun compensation by honeybees: How partially experienced bees estimate the sun’s course. Proceedings of the National Academy of Sciences, 91(10), 4471–4474. 10.1073/pnas.91.10.4471

Dyer, F. C., & Gould, J. L. (1981). Honey Bee Orientation: A Backup System for Cloudy Days. Science, 214(4524), 1041–1042. 10.1126/science.214.4524.1041

Fisher, N. I. (1993). Statistical Analysis of Circular Data (1st ed.). Cambridge University Press. 10.1017/CBO9780511564345

Freas, C. A., & Cheng, K. (2022). The Basis of Navigation Across Species. Annual Review of Psychology, 73(1), 217–241. 10.1146/annurev-psych-020821-111311

Freas, C. A., & Cheng, K. (2025). Diurnal Bull Ants Navigating by Moonlight: Polarised Light Homing in Myrmecia tarsata. Animal Behavior and Cognition. 10.1101/2025.02.21.639436

Freas, C. A., Narendra, A., & Cheng, K. (2017). Compass cues used by a nocturnal bull ant, Myrmecia midas. Journal of Experimental Biology, 220(9), 1578–1585. 10.1242/jeb.152967

Freas, C. A., Narendra, A., Lemesle, C., & Cheng, K. (2017). Polarized light use in the nocturnal bull ant, Myrmecia midas. Royal Society Open Science, 4(8), 170598. 10.1098/rsos.170598

Freas, C. A., Narendra, A., Murray, T., & Cheng, K. (2024). Polarised Moonlight Guides Nocturnal Bull Ants Home. 10.7554/eLife.97615.3

Freas, C. A., Wystrach, A., Narendra, A., & Cheng, K. (2018). The View from the Trees: Nocturnal Bull Ants, Myrmecia midas, Use the Surrounding Panorama While Descending from Trees. Frontiers in Psychology, 9. 10.3389/fpsyg.2018.00016

Islam, M., Deeti, S., Murray, T., & Cheng, K. (2022). What view information is most important in the homeward navigation of an Australian bull ant, Myrmecia midas? Journal of Comparative Physiology A, 208(5–6), 545–559. 10.1007/s00359-022-01565-y

Islam, M., Freas, C. A., & Cheng, K. (2020). Effect of large visual changes on the navigation of the nocturnal bull ant, Myrmecia midas | Animal Cognition. Animal Cognition, 23(6), 1071–1080.

Jander, R. (1957). Die optische Richtungsorientierung der Roten Waldameise (Formica Ruea L.). Zeitschrift für Vergleichende Physiologie, 40(2), 162–238. 10.1007/BF00297947

Klotz, J. H., & Reid, B. L. (1993). Nocturnal orientation in the black carpenter antCamponotus pennsylvanicus (DeGeer) (Hymenoptera: Formicidae). Insectes Sociaux, 40(1), 95–106. 10.1007/BF01338835

Lanfranconi, B. (1982). Kompassorientierung nach dem rotierenden Himmelsmuster bei der WüstenameiseCataglyphis bicolor. [PhD Thesis]. University of Zürich.

Massy, R., & Wotton, K. R. (2023). The efficiency of varying methods and degrees of time compensation for the solar azimuth. Biology Letters, 19(11), 20230355. 10.1098/rsbl.2023.0355

Mouritsen, H., & Frost, B. J. (2002). Virtual migration in tethered flying monarch butterflies reveals their orientation mechanisms. Proceedings of the National Academy of Sciences, 99(15), 10162–10166. 10.1073/pnas.152137299

Perez, S. M., Taylor, O. R., & Jander, R. (1997). A sun compass in monarch butterflies. Nature, 387(6628), 29–29. 10.1038/387029a0

Schmidt-Koenig, K., Ganzhorn, J. U., & Ranvaud, R. (1991). The Sun Compass. In P. Berthold (Ed.), Orientation in Birds (Vol. 60, pp. 1–15). Birkhäuser Basel. 10.1007/978-3-0348-7208-9_1

Spiecker, L., Laurien, M., Dammann, W., Franke, A., Clemmesen, C., & Gerlach, G. (2022). Juvenile Atlantic herring (Clupea harengus) use a time-compensated sun compass for orientation. Journal of Experimental Biology, 225(18), jeb244607. 10.1242/jeb.244607

Stalleicken, J., Mukhida, M., Labhart, T., Wehner, R., Frost, B., & Mouritsen, H. (2005). Do monarch butterflies use polarized skylight for migratory orientation? Journal of Experimental Biology, 208(12), 2399–2408. 10.1242/jeb.01613

Towne, W. F. (2008). Honeybees can learn the relationship between the solar ephemeris and a newly-experienced landscape. Journal of Experimental Biology, 211(23), 3737–3743. 10.1242/jeb.003640

Towne, W. F., & Moscrip, H. (2008). The connection between landscapes and the solar ephemeris in honeybees. Journal of Experimental Biology, 211(23), 3729–3736. 10.1242/jeb.022970

Ugolini, A., Galanti, G., & Mercatelli, L. (2013). Do sandhoppers use the skylight polarization as a compass cue? Animal Behaviour, 86(2), 427–434. 10.1016/j.anbehav.2013.05.037

Ugolini, A., Melis, C., & Innocenti, R. (1999). Moon orientation in adult and young sandhoppers. Journal of Comparative Physiology A: Sensory, Neural, and Behavioral Physiology, 184(1), 9–12. 10.1007/s003590050301

Warrant, E., & Dacke, M. (2016). Visual Navigation in Nocturnal Insects. Physiology, 31(3), 182–192. 10.1152/physiol.00046.2015

Wehner, R. (2020). Desert navigator: The journey of an ant. The Belknap Press of Harvard University Press.

Wehner, R., & Lanfranconi, B. (1981). What do the ants know about the rotation of the sky? Nature, 293(5835), 731–733. 10.1038/293731a0

Wehner, R., & Müller, M. (1993). How do ants acquire their celestial ephemeris function? Naturwissenschaften, 80(7), 331–333. 10.1007/BF01141909

Wystrach, A., Schwarz, S., Schultheiss, P., Baniel, A., & Cheng, K. (2014). Multiple sources of celestial compass information in the Central Australian desert ant Melophorus bagoti. Journal of Comparative Physiology A, 200(6), 591–601. 10.1007/s00359-014-0899-x

