## Supplemental Figure for "Nocturnal Navigation via a Time-Compensated Lunar Compass in Bull Ants"

Supplemental Figures

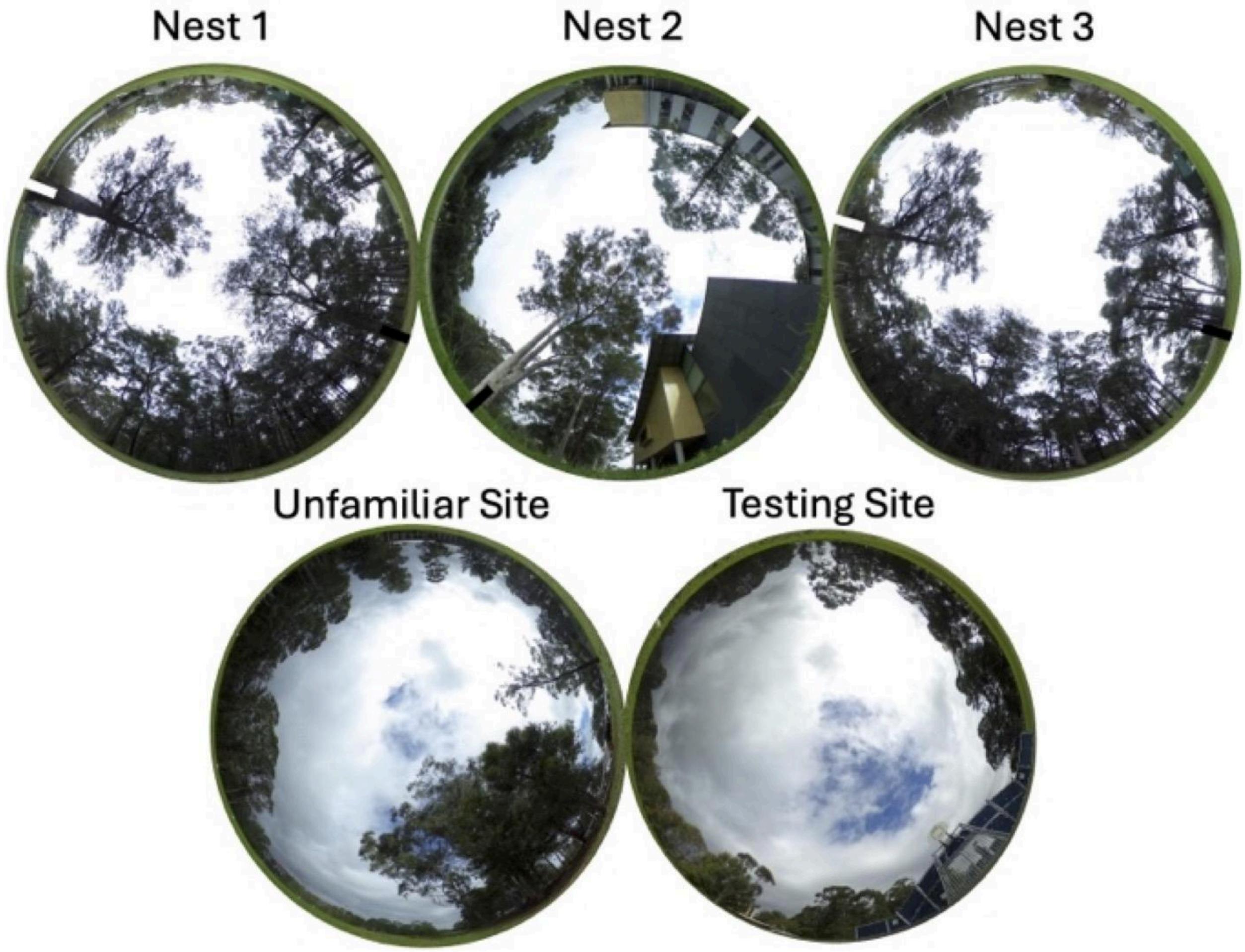


**Supplemental Figure 1.** Images of the landmarks, sky and canopy cover at nests and testing sites. Photos of nests were taken at the on-route midpoint between the foraging and nest trees. Black bars represent each nest tree while white bars represent the foraging tree. The unfamiliar site panorama/sky was used for the Unfamiliar condition, where foragers were held with a clear view of the moon but at a previously unfamiliar site. The Test site image was where all foragers were tested.

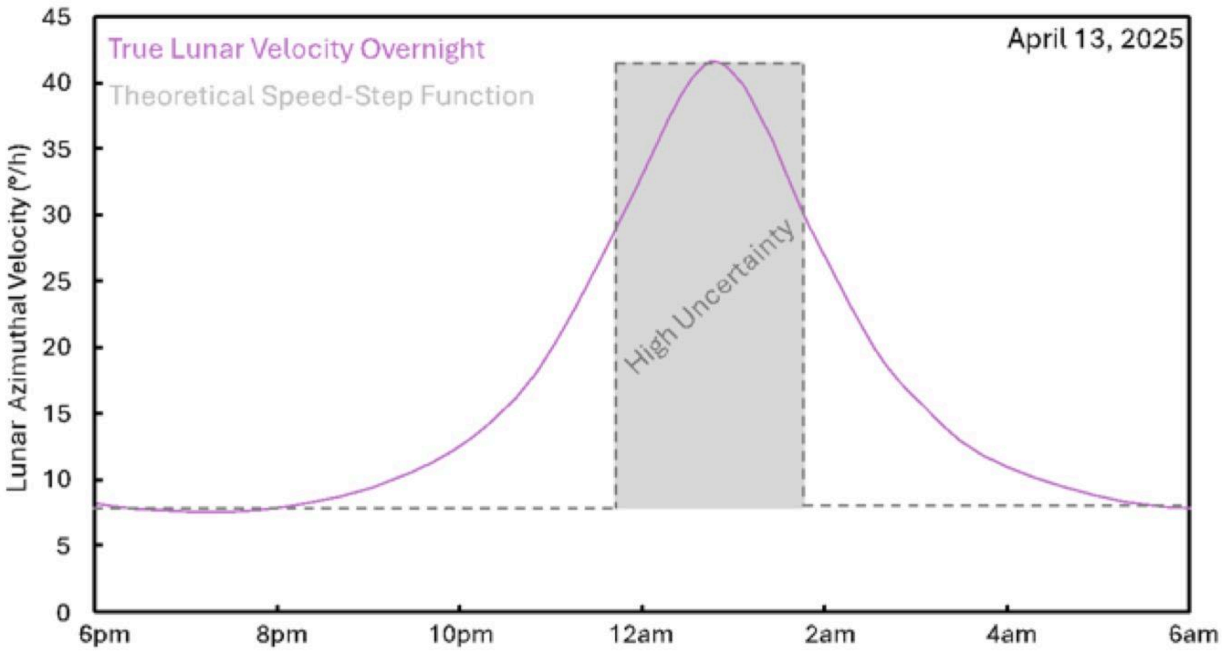


**Supplemental Figure 2.** Lunar Speed-Step-Function prediction for the lunar time compensated compass. The true lunar speed spike on the 13th of April 2025 (pink) overlayed with the proposed lunar speed-step-function in the nocturnal path integrator (grey). The grey area represents an area of temporal uncertainty within the system predicting this speed-step, which could explain which foragers underestimate lunar speed during this period, but remain well oriented to the setting moon, post speed-step.

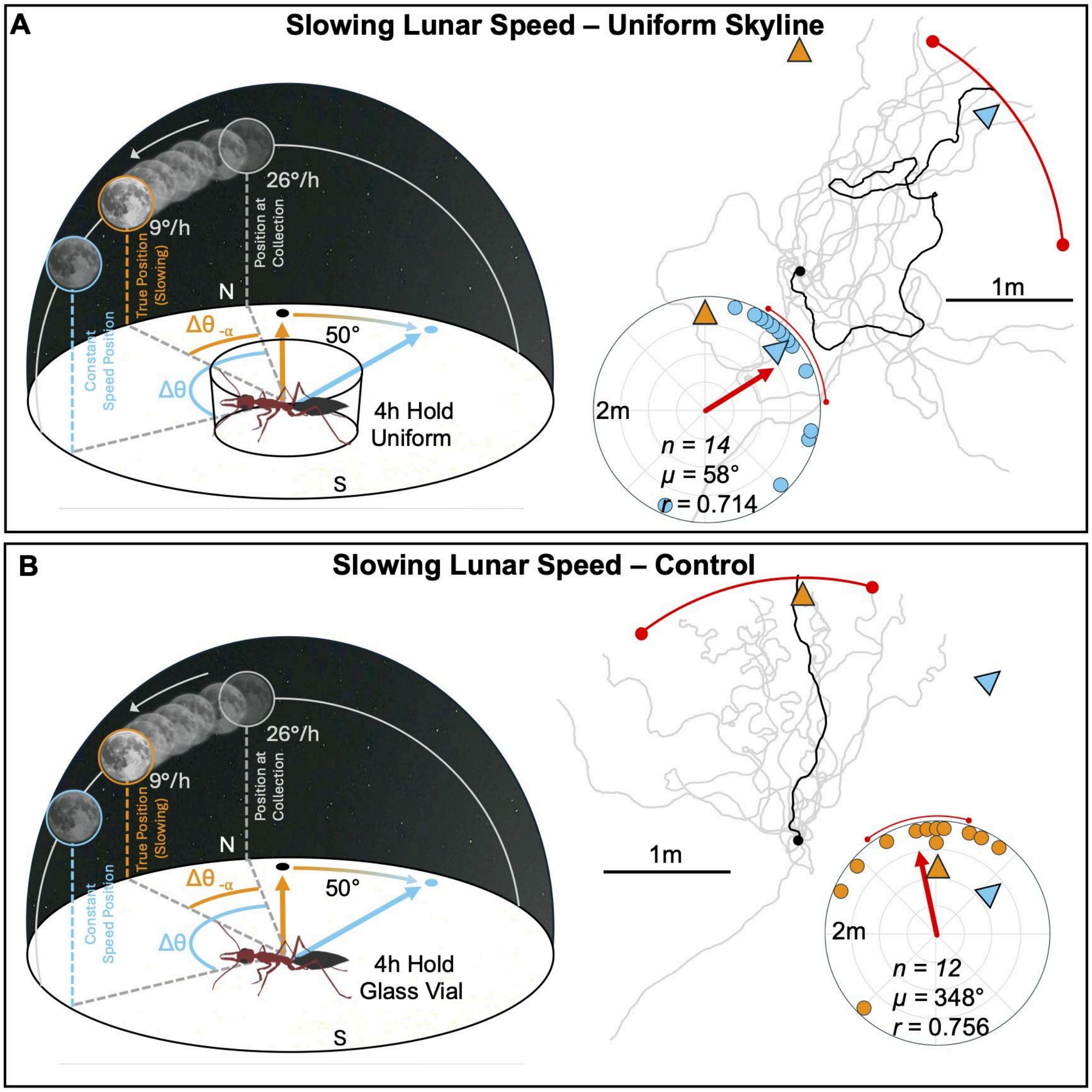


**Supplemental Figure 3.** Experimental Diagrams, Forager Paths and Circular Plots of Conditions and Forager Headings, at 2m from release, when allowed to observe a slowing moon with or without access to a directionally informative terrestrial skyline. Within the condition diagrams, colours represent different lunar predictions and the resulting predicted vector directions (triangles); with blue representing linear extrapolation of lunar azimuthal speed from last observation at collection, while orange represents the true deceleration of the lunar ephemeris. Foragers were allowed to collect a path integrator vector to their foraging tree under a fast moving moon near its apex, then either (A) held with a uniform skyline or (B) held in a glass vial while the moon decelerated during its descent and were then tested at a distant unfamiliar site. Red arrows within circular plots denote the length (r) and direction (μ) of the mean vector, while the red arc represents the 95% CI. n, number of individuals. 0° represents the true vector direction.
